# An α-π transition in S6 shapes the conformational cycle of the bacterial sodium channel NavAb

**DOI:** 10.1101/2022.06.21.496945

**Authors:** Koushik Choudhury, Rebecca J Howard, Lucie Delemotte

## Abstract

Voltage gated sodium channels play an important role in electrical signaling in excitable cells. In response to changes in membrane potential, they cycle between nonconducting and conducting conformations. With recent advances in structural biology, structures of sodium channels have been captured in several distinct conformations, thought to represent different functional states. However, it has been difficult to capture the intrinsically transient open state. We recently showed that a proposed open state of the bacterial sodium channel NavMs was not conductive, and that a conformational change involving a transition to a *π* helix in the pore-lining S6 helix converted this structure into a conducting state. However, the relevance of this structural feature in other sodium channels, and its implications for the broader gating cycle, remained unclear. Here, we propose a comparable open state of another class of bacterial channel from *Aliarcobacter butzleri* (NavAb), with characteristic pore hydration, ion permeation and drug binding properties. Furthermore, we show that a *π*-helix transition can lead to pore opening, and that such a conformational change blocks fenestrations in the inner helix bundle. We also discover that a region in the C-terminal domain can undergo a disordering transition proposed to be important for pore opening. These results support a role for a *π*-helix transition in the opening of NavAb, enabling new proposals for the structural annotation and drug modulation mechanisms in this important model sodium channel.

## Introduction

Voltage-gated sodium ion channels (Navs) are membrane proteins that selectively transport sodium ions across cell membranes. They open in response to membrane depolarization, and, in excitable cells, are responsible for the fast rising phase of the action potential. They are further characterized by rapid inactivation, during which the pore becomes non-conductive despite continued depolarized conditions (1). Eukaryotic Nav channels are pseudotetrameric single-chain proteins consisting of around 2000 residues. Bacterial sodium channels, on the other hand, have a simpler homotetrameric architecture, where each of four identical subunits comprises around 270 residues. Because bacterial channels share important aspects of structure and function with eukaryotic Nav channels, they have been used as prototypes to understand structure-function relationships in this superfamily of channels (2).

Several structures of bacterial (3-20) and eukaryotic Nav channels (21-27) have been determined. A prototypical Nav channel consists of three types of domains - voltage sensing domain (VSD), a pore domain and an intracellular C-terminal domain (CTD, Figure 1.A). Each VSD constitutes a four-helix bundle made of S1-S4, where S4 contains several arginine residues responsible for sensing the changes in potential difference across the membrane. Each channel contains four VSDs, each contributed by one subunit, that are arranged around a central pore. In their resting state, the VSDs’ S4 helices are stabilized in an ‘inward’ or ‘down’ position. Upon activation, the S4 helices transition into an ‘outward’ or ‘up’ position (7). The pore domain consists of a tetrameric inverse cone-shaped arrangement of the S5 and S6 helices from each of the four subunits. The pore is thought to open and close at the so-called intracellular bundle crossing, where an expanded pore likely represents an open state and a constricted pore likely corresponds to a closed state. In the resting state of the channel, the VSDs are in a resting conformation and the pore is non-conductive. In the open state, the VSDs are activated, and the pore is dilated. In the inactivated state, the VSDs remain activated, but the pore is non-conductive. In addition to these transmembrane domains, bacterial Nav channels possess a CTD that extends into the cytoplasm. This domain consists of two main regions - a neck region (proximal to the transmembrane region of the protein) and a distal coiled-coil region (Figure 1.A). Disordering of the neck region of the CTD favors activation and pore opening in bacterial sodium channels such as NavAe and NavSp (19, 20) The CTD is also involved in channel inactivation through an unknown mechanism (28).

**Figure 1:**
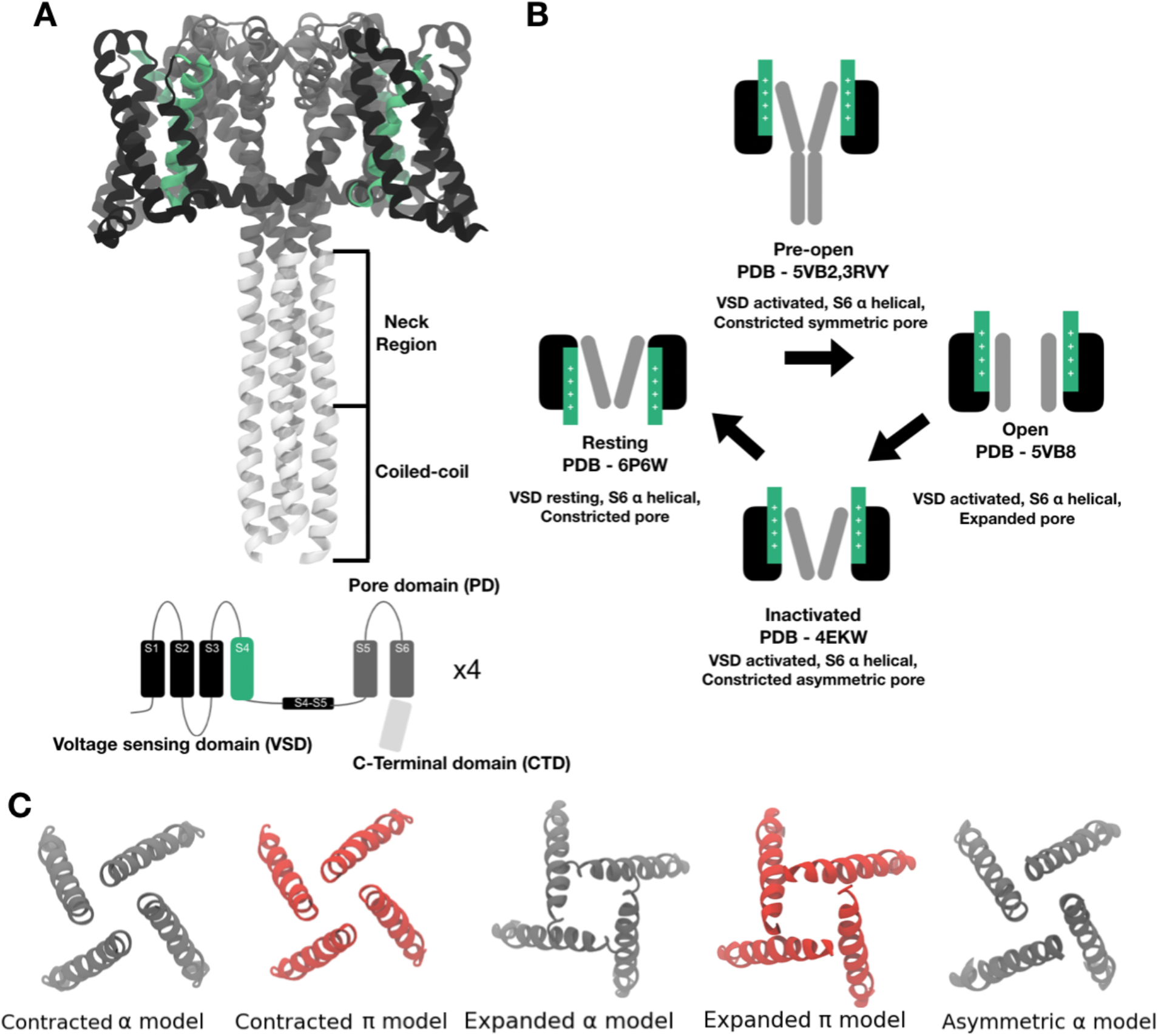
Structure and proposed functional cycle for NavAb. **A**. Full length structure of NavAb (PDB ID - 5VB2), initially described as a pre-open state, with activated voltage-sensor domains (VSD), contracted pore and ordered C-terminal domain (CTD). Inset shows a cartoon representation of one subunit highlighting the relevant domains - black - VSD, green - S4, dark grey - Pore domain and light grey - C-terminal domain (CTD) **B**. Currently proposed functional cycle for NavAb based on structures resolved in different functional states. **C**. Intracellular view of the pore conformation of the different models considered in this work; grey - α models; red - *π* models.

Structures of the sodium channel from *Aliarcobacter butzleri* (NavAb) have been described as representing three functional states (7,9,10). Based on these structures, the following functional cycle has been proposed (Figure 1B,C). Starting from a resting state (PDB ID - 6P6W) with VSDs ‘down’ and the pore in a constricted conformation, the channel transitions to a metastable pre-open state in which the VSDs are activated and the pore non conductive. This transition may occur in two steps, first to a state with a highly constricted and symmetric pore (PDB ID - 5VB2) followed by a second with a slightly wider and symmetric pore (PDB ID - 3RVY). After this, the pore transitions to an open state that features an expanded pore (PDB ID - 5VB8). Finally, the channel moves to an inactivated state wherein the VSDs are still activated but the pore is constricted and asymmetrical (C2 symmetry - PDB ID - 4EKW). In 5VB2, an ordered coiled-coiled CTD was resolved, but this domain is missing from all other structures.

Although this proposed cycle is consistent with some functional predictions, most structures of NavAb are of constructs containing one or more mutations, and it is thus unclear how well they represent functional states of wild-type (WT) NavAb. The open state has been especially difficult to capture, given the propensity of these channels to rapidly inactivate. Indeed, the structure of NavAb that is most likely to represent an open state, determined by x-ray crystallography, contains the pore-domain mutation I217C and is truncated after the S6 helix, thus deleting the CTD (10). As molecular dynamics simulations indicated that the pore of this mutant was hydrated and conducted sodium ions, this structure was assigned as an open state (10). However, functional properties of this structure upon reversion to the WT sequence remained unclear.

In our recent study of the related bacterial channel NavMs (29), we found that a structure initially thought to represent an open state, with an expanded pore with entirely α-helical S6 helices, was dehydrated and non-conductive to sodium ions. Introducing a short π-helical stretch in S6 in this channel resulted in a rotation of the S6 C-terminus and pore hydration, allowing sodium ion permeation in NavMs. In TRP channels, a π-helix formation and asparagine reorientation was recently proposed to be involved in pore opening (30-32). In addition, a reprocessing of the cryo-EM densities of Nav1.7 showed that the S6 helix in domain IV could be captured both in an α and in a *π*-state (33). Inspired by these observations, we hypothesized that the opening of NavAb may also require a transition to a π-helix in S6.

Local anesthetic drugs act by binding to sodium channels (34), several of them in a state-dependent manner. Bacterial sodium channels are known to be modulated by these drugs in a similar manner as their eukaryotic counterparts, making them useful tools to study state-dependent drug binding (2, 35). Depending on their protonation states, such drugs can access the Nav central cavity either via a hydrophobic pathway involving partitioning of the drugs in the membrane, or via a hydrophilic pathway across the intracellular ion gate (36). Indeed, structures have revealed the presence of fenestrations that connected the Nav central cavity to the membrane (9). These fenestrations were confirmed to play an important role in tonic block, whereby drugs access the central cavity when the pore is closed (5). The generation of a new open model poses the opportunity to study access of local anesthetics to their binding site in various states of NavAb.

In this study, we aimed to revise the proposed conformational cycle of NavAb, and to describe detailed features of the pore domain in putative resting, open and inactivated states of the channel. To do so, we first used molecular dynamics (MD) simulations to describe the full length structure of WT NavAb in a presumed open state. Reverting the I1217C mutation dehydrated the pore, leading to a model that was impermeable to sodium ions. Given our previous model of NavMs pore opening (29), we then constructed an alternate model of the NavAb pore, in which introducing a π-helix in S6 results in a hydrated and conductive pore. This open-pore model was validated by sodium ion and local-anesthetic permeation free energy calculations. We then used the open model to characterize state-dependent properties of hydrophobic fenestrations and lidocaine binding. Finally, we posited a new opening mechanism for NavAb, in which the formation of a π-helix in S6 and disordering of the neck region of the CTD are prerequisites for pore expansion.

## MATERIALS AND METHODS

### Model building

Four different NavAb structures were used in this study (Figure 1B,C) : a resting state (PDB ID 6P6W), in which the VSDs are at rest, the symmetric pore is contracted, and the CTD is truncated; a tentatively assigned open state (PDB ID 5VB8), in which the VSDs are activated, the symmetric pore is expanded, and the CTD is again truncated; and two decoupled states, tentatively assigned as pre-open (PDB ID 5VB2) and inactivated (PDB ID 4EKW) in which the VSDs are activated and the pore is contracted. In 5VB2, the pore domain has C4-symmetry and the CTD is ordered, while in 4EKW, the pore domain has only C2-symmetry (partly asymmetric) and the CTD is truncated. All the mutations in the structures were reverted to WT using CHARMM-GUI (37). Incomplete loops were also modeled using CHARMM-GUI (37).

Alternative models of NavAb with contracted or expanded pores were built by introducing a π-helix in a region of the S6 helix of 5VB2 and 5VB8, respectively. In all of these models, the VSDs remain activated. This was done using MODELLER 9.22 (38) following a protocol implemented in previous work on NavMs (29). Briefly, this consisted in shifting the structural alignment in S6 of the π-versus template α-models in order to introduce an unpaired backbone carbonyl. Specifically, we introduced one gap in the template (α) sequence prior to T1206, and a second in the target (π) sequence following D1219, thus shifting the region one position upstream from T1206 to the inner end of S6, and resulting in the formation of a π helix around T1206. This resulted in rotation of the pore helix following the π helix around T1206. As a result of this, residues facing the pore in the α-model rotate away from it in the π-model, while a different set of residues—including N1211—now face the pore [29]. Since all the residues in this region except N1211 are hydrophobic, the protonation states of the residues are not affected by the rotation.

For pore-opening Adiabatic Bias MD (ABMD, see below for details) simulations, we also built contracted-α and contracted-π NavAb models from PDB ID 5VB2 with the CTD deleted, otherwise following the procedure described above. This approach ensured that the protein length was consistent with the expanded-pore models.

### System preparation

Each model was embedded in a homogenous lipid bilayer consisting of 400 1-palmitoyl-2-oleoyl-*sn*-glycero-3-phosphocholine (POPC) using the CHARMM-GUI Membrane builder (37). Lidocaine-protein complexes were prepared by randomly placing the drug in the central cavity of the different models, respectively. Each system was hydrated by adding ∼45 Å layers of water to either side of the membrane. Lastly, the systems were neutralized with 150 mM NaCl. The Charmm36 force field was used to describe interactions between protein (39), lipids (40) and ions. In addition, we considered additional simulations using Charmm36m to study CTD disordering, as this particular forcefield is better suited to describe disordered proteins (41). Non-bonded fixes (NBFix) were considered in the description of interactions of sodium ions with carboxyl and carbonyl groups (42). The TIP3P model was used to describe all water molecules (43). Parameters for charged lidocaine were taken from our previous study (29).

### Simulation details

For the expanded-pore states (with or without lidocaine) and the protein-lidocaine system simulations, equilibration was done following the protocol reported in (10). Each system was minimized for 5000 steps using steepest descent and equilibrated with constant number of particles, pressure and temperature (NPT) for 30 ns, during which position restraints on the protein were gradually released. Initially the heavy atoms of protein were restrained with a force constant of 1000 kj/mol/nm^2^ for 10 ns. Then the protein backbone was restrained with a force constant of 1000 kj/mol/nm^2^ for 10 ns. Finally, the protein Cα atoms were restrained with a force constant of 1000 kj/mol/nm^2^ for 10 ns. For all the contracted models, equilibration was carried out in 6 steps according to the default CHARMM-GUI protocol, extending the duration of each step (First 3 steps were run for 2 ns each and the last 3 steps for 10 ns each) (37). During both equilibration and production, a time step of 2 fs was used, pressure was maintained at 1 bar through Berendsen pressure coupling, temperature was maintained at 300 K through Berendsen temperature coupling (44) with the protein, membrane and solvent coupled and the LINCs algorithm (45) was used to constrain the bonds involving hydrogen atoms. For long range interactions, periodic boundary conditions and particle mesh Ewald (PME) were used (46). For short range interactions, a cut-off of 12 Å was used. Production simulations were run with/without restraints for different systems as shown in Table 1, using Parinello-Rahman pressure coupling (47) and Nose-Hoover temperature coupling (48). Simulations were performed using GROMACS 2020.3 (49,50). Some simulations were performed under a transmembrane voltage of +/-750 mV, via application of an external electric field. The transmembrane voltage is calculated as the product of electric field and the simulation box length along which the field is applied. Such a transmembrane voltage did not cause any disruption to the protein/membrane over the relatively short simulation time considered.

**Table 1:** Summary table of equilibrium simulations performed. Restrained production simulations (where indicated) were used to prevent pore collapse in the absence of CTD.

| System | Equilibration | Production |
| --- | --- | --- |
| NavAb expanded α-model (5VB8) | 30 ns | 185 ns (unrestrained) |
| NavAb expanded α-model I217C (5VB8) | 30 ns | 250 ns (C-α restrained) |
| NavAb expanded α-model (5VB8) | 30 ns | 200 ns (C-α restrained) |
| NavAb expanded α-model (+750mV) | 30 ns | 200 ns (C-α restrained) |
| NavAb expanded α-model (-750 mV) | 30 ns | 200 ns (C-α restrained) |
| NavAb expanded π-model (5VB8) | 30 ns | 200 ns (C-α restrained) |
| NavAb expanded π-model (5VB8) | 30 ns | 154 ns (unrestrained) |
| NavAb expanded α-model + lidocaine | 30 ns | 200 ns (C-α restrained) |
| NavAb asymmetric $\alpha$ model + lidocaine | 30 ns | 200 ns (unrestrained) |
| NavAb resting $\alpha$ model + lidocaine | 30 ns | 200 ns (unrestrained) |
| NavAb expanded π-model + lidocaine | 30 ns | 200 ns (C-α restrained) |
| NavAb expanded π-model N1211L | 30 ns | 200 ns (C-α restrained) |
| NavAb contracted α-model (5VB2) with CTD - 300K<br>- Charmm36m | 36 ns | 200 ns (unrestrained) |
| NavAb contracted α-model (5VB2) with CTD - 320K<br>- Charmm36m | 36 ns | 308 ns (unrestrained) |
| NavAb contracted α-model (5VB2) with CTD - 360K<br>- Charmm36m | 36 ns | 309 ns (unrestrained) |
| NavAb contracted α-model (5VB2) with CTD - 400K<br>- Charmm36m | 36 ns | 215 ns, 223 ns, 232 ns<br>(unrestrained) |
| NavAb contracted π-model (5VB2) with CTD | 36 ns | 200 ns (unrestrained) |
| NavAb contracted π-model (5VB2 WT) without<br>CTD | 36 ns | Equilibrated structure used<br>for ABMD simulation $\times 3$ |
| NavAb contracted α-model (5VB2) without CTD | 36 ns | Equilibrated structure used<br>for ABMD simulation $\times 3$ |
| NavAb contracted α-model (5VB2 N1211L) without<br>CTD | 36 ns | Equilibrated structure used<br>for ABMD simulation $\times 3$ |

**Table 2:** Summary of the enhanced sampling free energy calculation performed.

| System | Enhanced Sampling technique | Production |
| --- | --- | --- |
| NavAb (expanded α-model) - Na<sup>+</sup> | AWH | 200 ns × 6 walkers |
| NavAb (expanded π-model) - Na<sup>+</sup> | AWH | 200 ns × 6 walkers |
| NavAb (expanded α-model) - lidocaine | AWH | 200 ns × 6 walkers |
| NavAb (resting $\alpha$ model) - lidocaine | AWH | 200 ns × 6 walkers |
| NavAb (asymmetric $\alpha$ model) - lidocaine | AWH | 200 ns × 6 walkers |
| NavAb (expanded π-model) - lidocaine | AWH | 200 ns × 6 walkers |
| NavAb expanded π-model (N214L) - Na<sup>+</sup> | AWH | 139 ns × 6 walkers |
| NavAb (resting $\alpha$ model) - lidocaine | Well-tempered metadynamics | 94 ns × 12 walkers |
| NavAb (asymmetric $\alpha$ model) - lidocaine | Well-tempered metadynamics | 94 ns × 12 walkers |
| NavAb (expanded π-model) - lidocaine | Well-tempered metadynamics | 140 ns × 16 walkers |
| NavAb (expanded α-model) - lidocaine | Well-tempered metadynamics | 140 ns × 16 walkers |
| Pore opening starting from contracted π-model (5VB2 WT) without CTD | ABMD | 840 ps × 3 replicas |
| Pore opening starting from contracted α-model (5VB2) without CTD | ABMD | 840 ps × 3 replicas |
| Pore opening starting from contracted π-model (5VB2 N1211L) without CTD | ABMD | 840 ps × 3 replicas |

### Ion permeation and drug permeation free energy calculations

The free energy profiles of sodium ions and lidocaine permeation along the pore axis were calculated using the accelerated weight histogram (AWH) method (51). In brief, for each equilibrated structure, we applied one independent AWH bias and simulated for 200 ns each with 6 walkers sharing bias data and contributing to the same target distribution. The target distribution chosen was a flat (uniform) distribution. Each bias acts on the z-axis defined using the center-of-mass *z*-distance between one central sodium ion/lidocaine (all atoms) and all atoms of S1178 residues in NavAb, with a sampling interval corresponding to the entire box length. The rate of change of each AWH bias was initialized by setting the initial average free energy error to 20 kj/mol and the diffusion constant was set to 0.00005 nm^2^/ps. To keep the solute close to the pore axis, the coordinate radial distance was restrained to stay below 10 Å by adding a flat-bottom umbrella potential. All the Cα atoms of the protein were restrained by imposing harmonic potentials with force constants of 1000 kJ mol^-1^ nm^-2^ to remain as close as possible to the experimental structures.The average potential of mean force (PMF) profile and the error bars were calculated from a single walker since the walkers communicate with each other. For all systems except the expanded π-model with lidocaine, the average PMF profile was calculated by taking the data from the last 100 ns in intervals of 10 ns and the error bars were calculated by calculating the standard deviation. For the expanded π-model with lidocaine, data from the last 50 ns at 10 ns intervals were chosen for calculating the average and standard deviation.

Additionally, the lidocaine permeation free energy along the pore axis was also calculated using multiple walker well-tempered metadynamics (52,53) in GROMACS 2020.5 patched with PLUMED 2.7.3 (54,55). Each bias acts on the z-axis defined using the center-of-mass *z*-distance between lidocaine (all atoms) and all atoms of S1178 residues in NavAb. The hills height and width was set at 1.2 kJ/mol and 0.05 nm respectively. Upper and lower restraints (as shown in the corresponding x-axes of the free energy plots) were applied to restrict the sampling space. For the models with constricted pores, the sampling was restricted to the central cavity of the pore. To keep the drug close to the pore axis, the coordinate radial distance was restrained to stay below 10 Å by restraining the x and y component of center-of-mass distance. The free energy profile was estimated using the sum_hills tool in PLUMED. The average free energy profile was calculated by taking the data from the last 30 ns in intervals of 5-10 ns and the error bars were estimated by calculating the standard deviation.

### Adiabatic biased molecular dynamics for pore opening

The pore opening was studied by performing Adiabatic Biased Molecular Dynamics (ABMD) (56) in GROMACS 2020.5 patched with PLUMED 2.7.3 (54,55). ABMD is a simulation method in which whenever the system moves closer toward the target system along the collective variable, the harmonic potential is moved to this new position, resulting in pushing the system toward the target value of the collective variable. The bias potential was applied along the distance between the center of masses of Cα atoms of I1216/I1217 and I1119 (for the α model and the π model, respectively). The force constant for the harmonic potential was set at 5000 kJ/mol/nm^2^ while the target value for the collective variable was set at 1 nm. The simulations were started from a contracted α and π model and were stopped when the system reached the expanded states. ABMD also outputs the square of force due to the bias potential. We calculated the total force by summing the forces applied along time. The first 840 ps was for each system and the replica was used for force calculation. Three replicas each were run starting from contracted α and π models respectively. The average total force across replicas was calculated and the error bars were estimated by calculating the standard deviation.

### Analysis

The water number density, water hydration free energy and pore radius were calculated using the channel annotation package (CHAP) (57). For the water density analysis, frames were extracted every 100 ps from the trajectories of the respective models. For the fenestration radius analysis, 200 equally spaced frames were extracted from the trajectories of the expanded/contracted α and π models. We calculated the time-averaged radius for each fenestration in each model. Since all the models are homotetrameric, we then calculated the average of the four fenestration radius and the uncertainty on the average estimate, as the average of the standard deviation, by taking the square root of the average of the squares of the standard deviations. Representative binding poses of lidocaine were obtained by performing clustering on frames extracted from the AWH calculations corresponding to free energy minima. Clustering was done using gmx cluster, with a cutoff of 0.1 nm. The center of the most populated cluster was then chosen as the representative binding pose. A similar approach was used to extract the most representative structures from the pore opening free energy profiles. Hydrogen bonding between N1211 and F1207 was calculated using gmx hbond. The rmsd for N1211 was calculated by first aligning the structures on their backbone atoms using gmx rms. The files necessary to reproduce the data herein can be found at https://osf.io/u8fg9/

## RESULTS

### The expanded-α structure of NavAb is non-conductive

To test the relevance to WT channel function of the proposed open-state structure of NavAb, in which the VSDs were captured in an activated state and the pore in an expanded conformation (PDB - 5VB8), we first reverted the I1217C mutations and carried out equilibrium MD simulations, still in absence of the CTD. Because of the α-helical structure of S6, and wide diameter of the pore, we call this model “expanded α”. In previous unrestrained simulations of this model, despite modest rotation of the pore-lining helix relative to the contracted ? model, the pore spontaneously collapsed to a dehydrated and nonconducting conformation (10,58). We therefore restrained the C-α positions to maintain the pore in an expanded state. Analyzing the pore hydration by calculating the time-averaged water number density along the central axis showed that the pore is nonetheless dehydrated close to the activation gate (-1 nm along the pore axis, Figure 2A, S1A), likely due to hydrophobic residues (V1213) lining the pore (Figure 2C), and in contrast to the I1217C mutant (Figure S1D).

**Figure 2:**
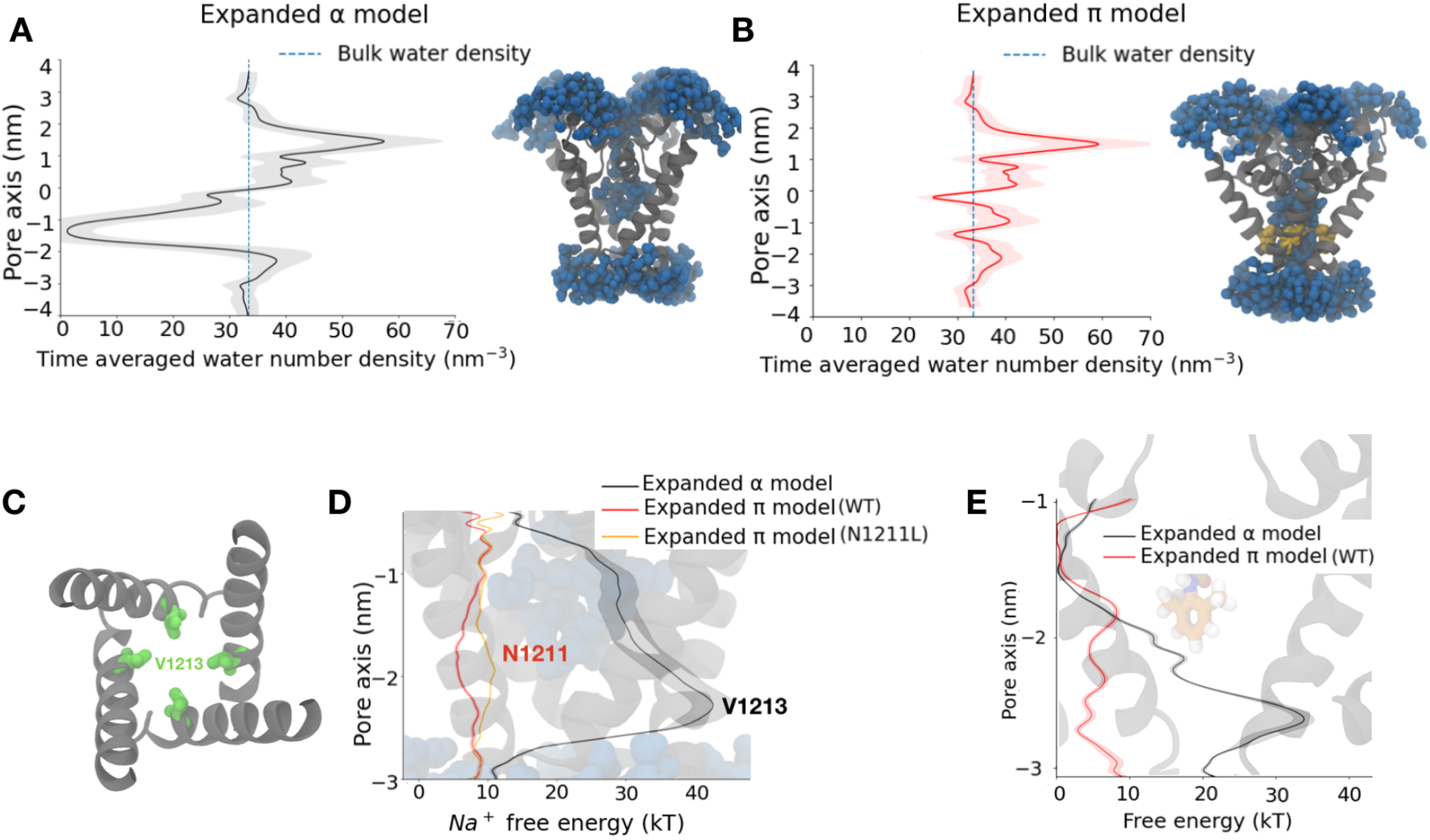
The *π* model is a functional open state based on pore hydration, and on sodium ion and lidocaine permeation free energy profiles. **A**. Time averaged water number density along the central pore axis of the expanded α model. Standard deviation shown in gray shading, the blue dashed line shows the bulk water density and the inset shows a snapshot from the expanded α model simulation wherein the pore is dehydrated close to the activation gate (grey helices - protein; blue spheres - water molecules). **B**. Time averaged water number density along the central pore axis of the expanded *π* model. The inset shows a snapshot from the expanded *π* model simulation with a completely hydrated pore (grey helices - protein; blue spheres - water molecules; yellow sticks - conserved asparagine). **C**. Hydrophobic constriction at V1213 (green) leading to pore dehydration in the expanded α model (view from the extracellular medium). **D**. Sodium ion permeation free energy profile for the expanded α model (black), and for WT (red) and N1211L (orange) variants of the expanded *π* model along the central pore axis of the lower half of the S6 helix. **E**. Lidocaine permeation free energy profiles for the expanded α model (black) and expanded *π* model WT (red) along the central pore axis of the lower half of the S6 helix. Cartoon representations in the background of D and E show the lower half of the S6 helix.

We next hypothesized that a transmembrane potential difference could affect pore hydration. Previous studies have indeed shown that applying external electric fields can promote hydration of hydrophobic pores (59). For the expanded-α model of NavAb, an external transmembrane voltage of -750 mV did not change pore hydration substantially. Applying a transmembrane voltage of +750mV increased hydration, but with a final water density still lower than that of bulk water (Figure S2). Moreover, the pore was only transiently hydrated, alternating between wetted and dewetted states (Figure S1B,C).

### Introducing a π-helix in S6 results in a conductive channel

Inspired by our previous work on NavMs (29), we next hypothesized that a π-helix in S6 might stabilize a conducting state of NavAb. Introducing this modification (see Methods), we produced an “expanded π” model with activated VSDs which we interrogated using MD simulations. Simulating this model resulted in a consistently hydrated pore (Figure 2B), in contrast to the expanded-α model (Figure S2). To further confirm that the expanded-π model represents a functional open state, we calculated sodium ion and lidocaine permeation free energy profiles across the channel gate using enhanced sampling. There was a substantial barrier to sodium (∼ 40 kT) and lidocaine (∼ 20 kT) exit from the central cavity in the expanded-α model, while there was no barrier in the expanded-π model (Figure 2 D,E, Figure S3-4,S7). To simplify comparisons to the expanded α model, we similarly performed these runs in the absence of the CTD and with Cα atoms restrained; without restraints, backbone motions still led to pore dehydration, consistent with a role for the CTD in stabilizing the expanded backbone configuration (Figure S8).

The introduction of this π-helix resulted in the rotation of a conserved asparagine, N1211, towards the NavAb pore lumen (Figure 2B). Mutating this pore-facing asparagine to a hydrophobic residue (N1211L) did not affect pore hydration (Figure S2, S5), but slightly increased the barrier to sodium-ion permeation at the site of mutation (Figure 2D). N1211 also likely plays an important role in stabilizing the π-helix by forming a hydrogen bond with the backbone carbonyl of F1207. Indeed, N1211 is less mobile in the expanded-π model than in the expanded-α model (Figure S6). Taken together, these results are consistent with the expanded-π model representing a functional open state, and with N1211 possibly contributing to sodium-ion permeation.

### Local anesthetic binding poses and access pathways vary with functional state

We hypothesized that a structural transition in S6 may impact the accessibility of neutral drugs to their binding site in the pore cavity through the pore-lining fenestrations. To test this, we calculated the time-averaged radius of the four fenestrations in the contracted-α, contracted-*π*, expanded-α and expanded-*π* models of NavAb. Comparing the fenestration radii for these models, we found that the *π* models appear to have narrower fenestrations than the α models (Figure 3A,B). Indeed, the fenestrations are obstructed by large hydrophobic residues F1207 and I1210 in both the contracted- and expanded-*π* models (Figure 3B). This observation indicates that hydrophobic drugs can access the central cavity only in the context of a fully α-helical S6, irrespective of pore diameter.

**Figure 3:**
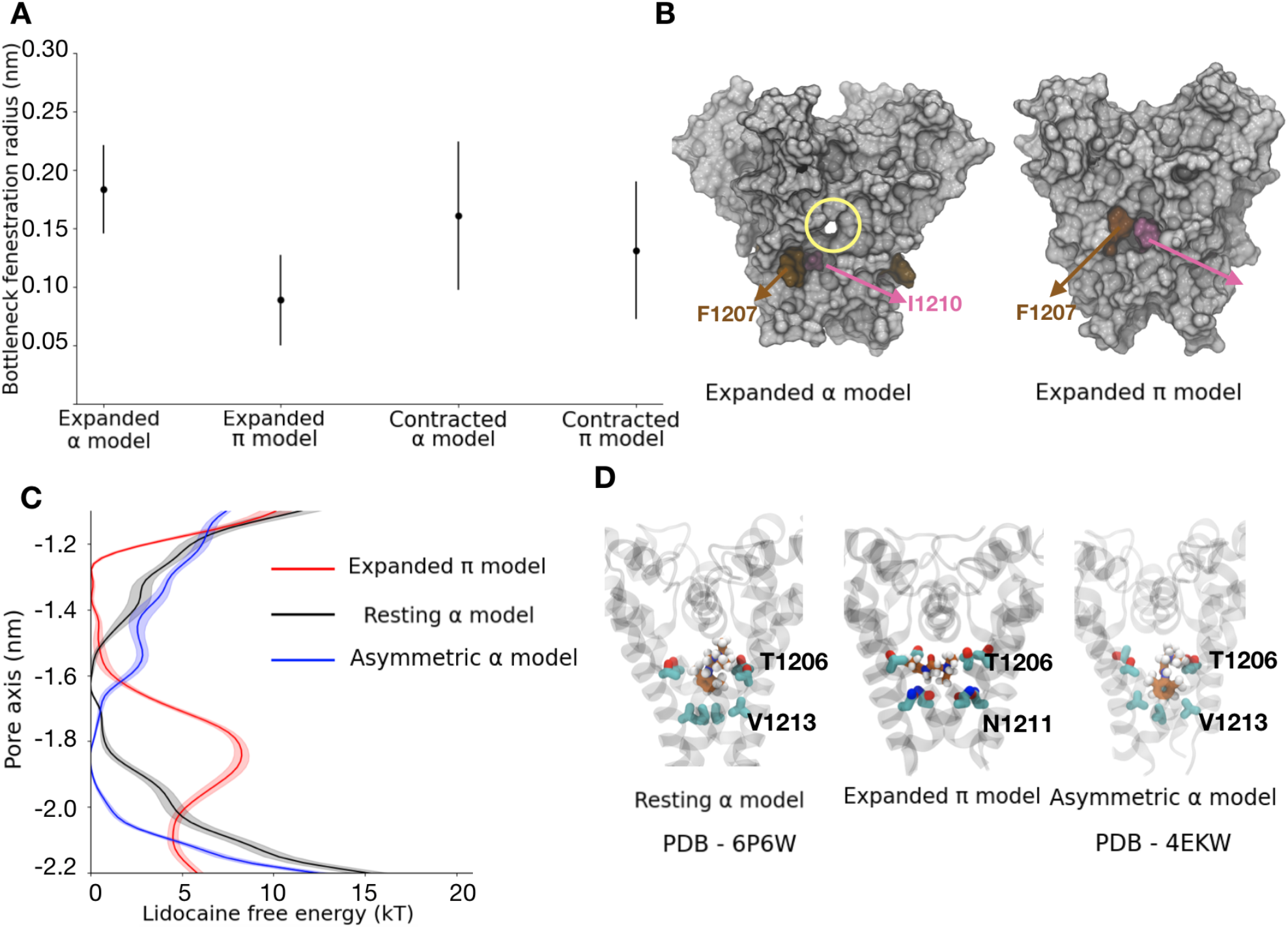
Relationship between fenestrations and *π*-helix formation. **A**. Time and subunit averaged radius of fenestrations for different models. For each model, we plot the average bottleneck radius across four subunits and the error bars represent the average standard deviation. **B**. Surface representation showing the open (yellow circle) and closed fenestrations in the expanded α and *π* models respectively. The fenestrations in the *π* models are closed by F1207 (brown) and I1210 (pink) residues. **C**. Lidocaine permeation free energy profiles for the resting model (black), expanded *π* model (red) and asymmetric α model (blue) along the central pore axis of the lower half of the S6 helix. **D**. Lidocaine binding site in the resting model, expanded *π* model and asymmetric α model. Lidocaine (sticks, carbon atoms in orange) is bound closer to the selectivity filter in the resting model and expanded *π* model, with a common binding site at T1206 (sticks, carbon atoms in cyan). In the asymmetric α model, the drug binding site has shifted more towards the intracellular end away from the selectivity filter.

We next investigated state-dependent drug access to the central cavity via the hydrophilic pathway by calculating lidocaine permeation free energies across the channel gate in different functional states of NavAb. We particularly compared our presumed-open expanded-*π* model to two previously reported structures, one with resting VSDs and a symmetrically contracted pore (assigned resting, PDB ID 6P6W), and one with activated VSDs and an asymmetric conformation of the pore (assigned inactivated, PDB ID 4EKW). Unlike the expanded-*π* model, the resting and inactivated models featured high barriers to drug access from the intracellular side (Figure 3C, Figure S10-11). In all three models, free-energy profiles indicated that the drug binds in the central cavity close to the selectivity filter, forming contacts with T1206 in both the resting and the expanded-*π* models. However, in the resting model, lidocaine was oriented parallel to the pore axis with its charged group pointing towards the selectivity filter, while in the expanded-*π* model it was oriented parallel to the membrane plane (Figure 3D). This difference could be due to the presence in the expanded-*π* model of the hydrophilic sidechain of N1211, which accommodates the charged group of lidocaine. Note however that lidocaine binding does not affect the flexibility of N1211 (Figure S6). In the asymmetric inactivated model, the drug was oriented perpendicular to the membrane plane and shifted further towards the intracellular end of the pore, closer to V1213 (Figure 3D). These results indicate that there is a change in the binding pose of lidocaine as the channel transitions between different functional states, and that the drug can access its binding site via the intracellular gate exclusively in the *π* model.

### Neck region of the C-terminal domain can undergo disordering

Experimental structures and electrophysiology recordings have indicated a disordered segment in the neck region of the CTD in certain states of several bacterial sodium channels (20). This conformational change is proposed to be important for pore opening. To test whether this region can undergo disordering in NavAb, we subjected the full-length constricted-α structure of NavAb (PDB ID - 5VB2) to unrestrained simulations at elevated temperatures (300K, 340K, 360K and 400K, the latter in triplicate). Simulations at 320K and 360K showed limited disordering in the CTD (figure 4B,C) while those at 400K showed a complete disordering of the neck region (Figure 4D) spanning residues Q1226 to I1243 (figure 4D). The secondary structure was overall preserved in the rest of the protein at 400K (Figure 4D), suggesting that the CTD neck is specifically prone to losing its secondary structure.

**Figure 4:**
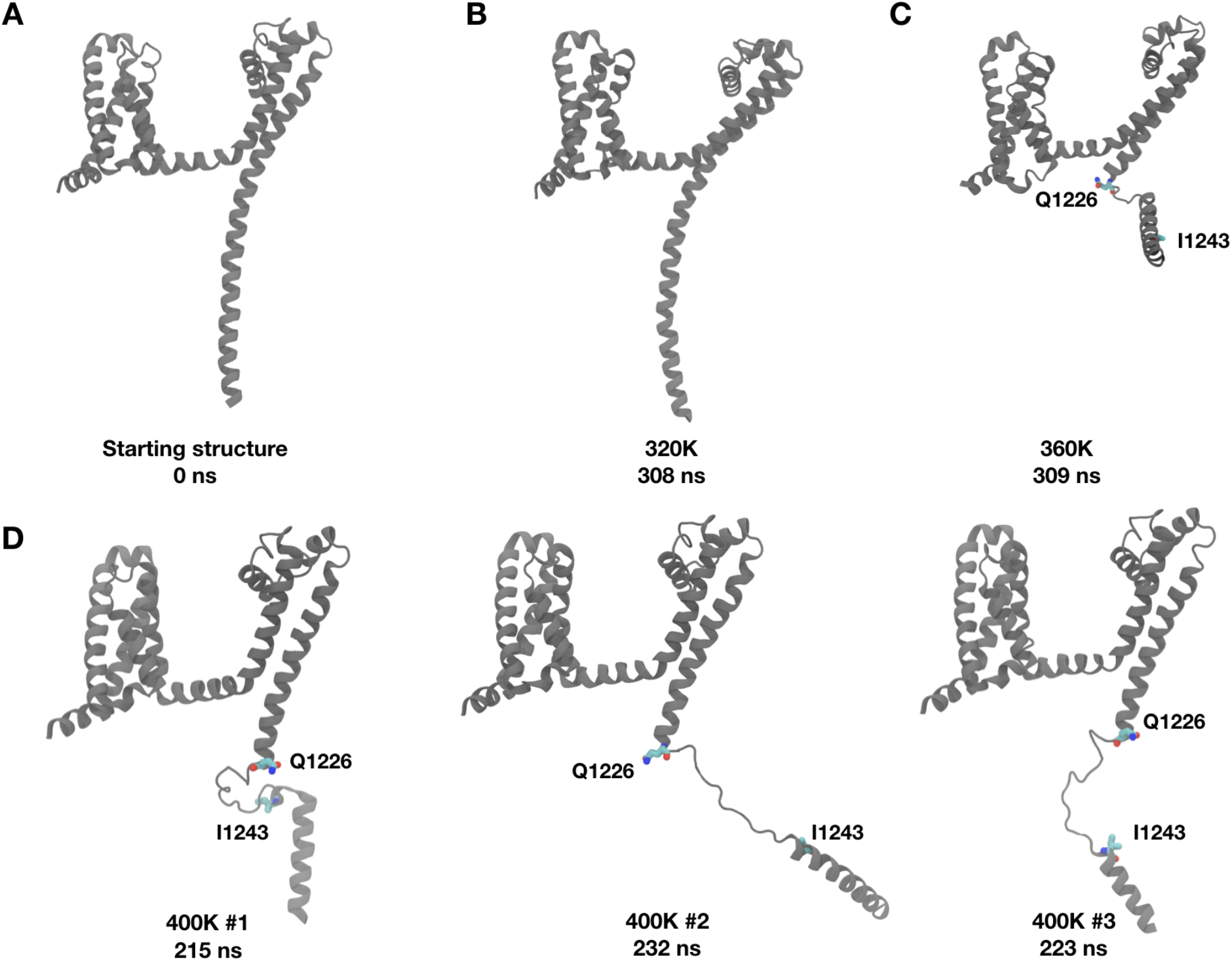
The neck region of the CTD proximal to the transmembrane helices can disorder. **A**. Cartoon representation of one subunit of the starting structure. **B**. Cartoon representation of one subunit of the final structure after 308 ns simulation at 320K **C**. Cartoon representation of one subunit the final structure after 309 ns simulation at 360K, showing disordering of the neck region. **D**. Cartoon representations of one subunit of the final structure after simulating each replica at 400K. The neck region between Q1226 and I1243 displays disordering.

### A *π*-helix transition results in pore opening

The results above suggested that a *π*-helix in S6, along with an expanded pore, results in a open state of NavAb. In contrast, a contracted and entirely α-helical S6, as resolved in the closed experimental structure, results in an obstructed pore. Pore opening therefore involves an α-to *π*-helix transition in S6 on top of pore expansion. To determine whether the α-to *π*-helix transition occurs before or after expansion of the pore, we characterized the force needed to open the pore in presence of an α- and of a *π*-helix. In the contracted-α model, N1211 is pointing away from the pore into the peripheral cavity, attracting water from the cytoplasm into peripheral cavities (Figure 5A). We propose that these water-filled peripheral cavities, as well as the attractive interaction between hydrophobic pore-facing residues at the activation gate, could impede pore opening. In the contracted-*π* model, on the other hand, N1211 faces the pore lumen rather than the peripheral cavities, which are hence no longer hydrated (Figure 5B). These observations could constitute a rationale for the low propensity of the α model to expand.

**Figure 5:**
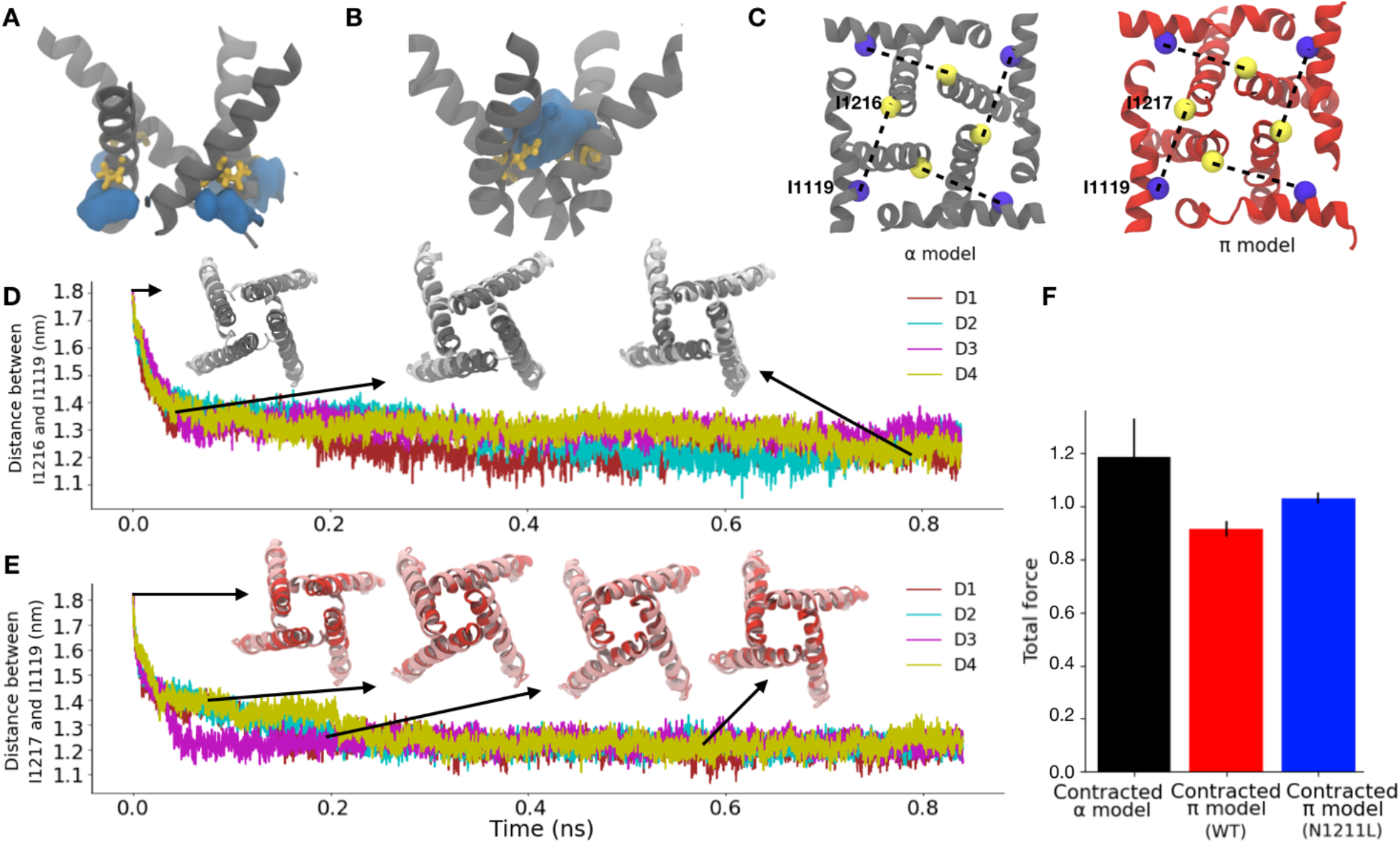
A *π* helix facilitates pore opening. **A**. N1211 (yellow sticks) pointing away from the central pore and hydrated peripheral cavities (water-filled volume represented as a blue surface) in the contracted α model. **B**. N1211 (yellow sticks) pointing into the hydrated pore (water-filled volume represented as a blue surface) in the contracted pi model. **C**. Distances (dashed lines) from I1119 (blue) to either I1216 or I1217 (yellow) in the α-model (left) or π-model (right) respectively, used as collective variables applied to pore opening in ABMD simulations. **D**. Representative replica of the time evolution of the pore opening ABMD simulation starting from the contracted α model. Pore configurations at different time points of the simulation are shown in inset as dark gray ribbons, overlaid with the expanded α model represented in silver. **E**. Representative replica of the time evolution of the pore opening ABMD simulation starting from the contracted *π* model. Pore configurations at different time points of the simulation are shown in inset as red ribbons, overlaid with the expanded *π* model represented in pink. **F**. Bar plot reporting the average total force required for pore opening starting from a contracted α model (black), a WT contracted *π* model (red) and a N1211L mutant contracted *π* model (Blue)

To add further quantitative support to this rationale, we performed ABMD simulations to open the pore starting from contracted *π* and α model, using as collective variable the distance between Cα atoms of I1216 (α model)/I1217 (*π* model) and I1119 across the four subunits (Figure 5C). The collective variable was chosen because these residues are located at the intracellular ends of S6 and the S4–S5 linker, respectively, such that the distance between them falls dramatically during transition from the pre-open/contracted to open states of the activated channel. This collective variable indeed enabled sampling of expanded-pore conformations from simulations initiated from contracted pores (Figure 5D, 5E, S16-18) and did not lead to substantial unwinding or other nonphysical transitions (Figure 5D, 5E, insets). We performed three replicates, each starting from the contracted *π* and α models, and calculated the average total force across the replicates. The average total force for pore opening was lower when the simulations were initiated from contracted *π* model relative to contracted α model, indicating that the *π* model is more likely to open than the α one (Figure 5F). To further characterize whether the presence of the conserved Asn contributes to lowering the energy needed to open the *π* model, we repeated the same simulations in the presence of the N1211L mutation, thus removing the only hydrophilic residue in this region. The average total force for pore opening increased in N1211L relative to the WT *π* model (Figure 5D), indicating that, indeed, N1211 contributes to a favorable opening of the pore, and possibly rationalizing its conservation across Nav channels.

## Discussion

Our first observation in this work was that the expanded-α experimental structure of NavAb does not convincingly model a functional open state. Indeed, in simulations of this model, the pore was dehydrated, preventing sodium ion permeation across the gate and lidocaine access to its binding site in the central cavity. However, introducing a short *π*-helix in S6 produced a consistently hydrated pore which allowed sodium/lidocaine permeation. One contributor to this phenomenon is likely the conserved asparagine (N1211), where mutation to leucine slightly increased the free-energy barrier to ion permeation.

What is then the mechanism of transition between the resting/closed and the activated/open states? We found that the CTD of NavAb has a region proximal to the membrane that can undergo a disordering transition. As similar disordering is thought to facilitate pore opening in other bacterial channels, a similar mechanism may contribute to opening in NavAb. Indeed, our understanding of the involvement of the CTD in the gating mechanism had been hampered by the lack of resolution in that domain in most structures of NavAb. Conducting simulations at high temperatures revealed a propensity for the neck region to disorder, in agreement with observations in other bacterial channels (20). Although not the focus of this study, future experimental or computational work — for example, temperature-based replica exchange simulations — could give us insights into the free energy needed to unfold this region in the various pore states, contributing to our understanding of the role played by this region.

Our pore-opening studies further showed that it is unfavorable to expand a constricted pore with an α-helical S6. Thus, channel activation could include a transition from α to *π*-helix in S6, via a state comparable to our contracted-*π* model. Indeed, the eukaryotic channel NavPas was recently captured in a conformation consistent with such a pre-open functional assignment, with activated VSDs, a contracted pore, *π* helices in all of its S6 segments, and a dissociated inactivation particle (60,61). The symmetric expanded-α model of NavAb, which was initially assigned as open (10), may instead represent an inactivated state of the WT channel, possibly contributing to a multi-step inactivation process along with the asymmetric contracted-α structure (9). Resolving this complex inactivation mechanism thus remains an area of active research, since the order in which *π* to α transitions, pore contraction, and possible changes in symmetry occur is currently unknown. It has been observed in TRPV1 channels that a conformation with a hydrated pore usually features dehydrated peripheral cavities between the S6 and S4-S5 helices, whereas conformations with a dehydrated pore have hydrated peripheral cavities (30). In the NavAb contracted-α model, N1211 attracts water into the peripheral cavities, apparently impeding pore opening. In the contracted-*π* model on the other hand, N1211 faces the pore lumen and stabilizing hydrophobic interactions at the gate are no longer present. This remodeling appears to favor opening of the hydrophobic activation gate, as witnessed in the pore-opening ABMD simulation of the *π* model and of the N1211L mutant.

Putting together these results, we propose a modified mechanistic picture for the functional cycle of bacterial sodium channels (Figure 6). Five important processes occur during the transition between the resting and open state - the VSDs activate, an α to *π* transition occurs in S6 resulting in rotation of the inner half of the helix, peripheral cavities dehydrate, the CTD disorders and the pore expands. Activation of the VSDs in response to membrane depolarization is presumed to be the initiating step in this cycle. Then, we propose that the α to *π* transition occurs, resulting in dehydration of peripheral cavities, finally followed by pore expansion. We also propose that the CTD disorders during the opening transition, though our data do not definitively indicate at what stage this occurs. We hypothesize that the energy to overcome barriers associated with *π*-helix formation, CTD disordering and pore opening, is contributed at least in part by coupling to VSD activation, which is favorable thanks to membrane depolarization. Nevertheless, as the mechanism of VSD-pore coupling in Nav channels is still being investigated by many groups (62,63,64), it is currently challenging to devise a quantitative mechanism.

**Figure 6:**
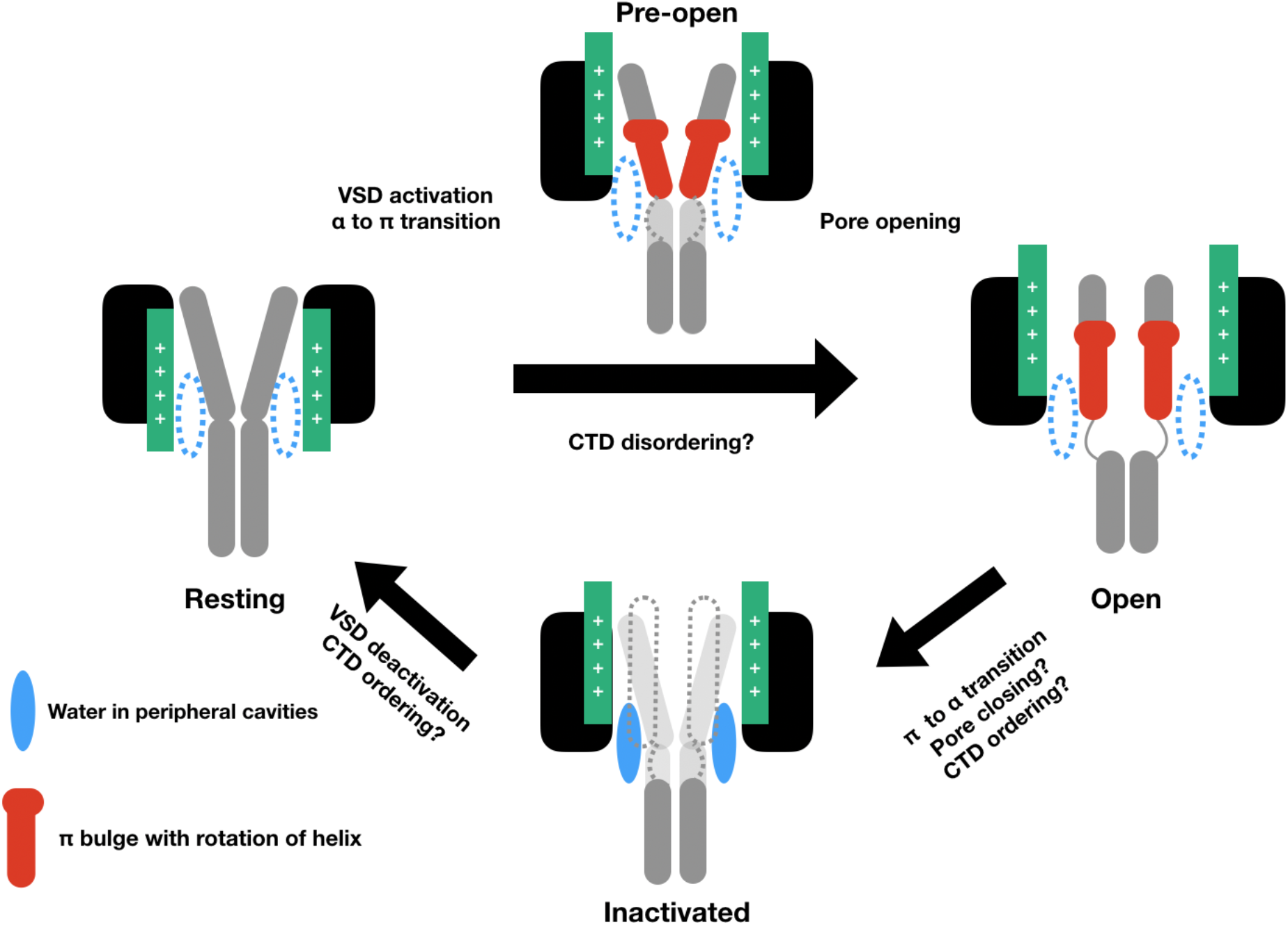
Proposed functional cycle for NavAb. Starting from the resting state with α helical S6, NavAb transitions to a pre-open state with VSDs activated and α to *π* transition. This conformation allows for favorable pore opening. Pore opening also involves CTD disordering, although the point at which this happens remains unclear. Inactivation involves transition from *π* to α, but whether the pore is expanded, contracted in a symmetric or asymmetric manner, or even transitions between different conformations, remains unclear. Return to the resting state involves deactivation of the voltage-sensor domains.

Finally, we considered what aspects of the proposed mechanism for bacterial-Nav gating and inactivation may be applicable to eukaryotic channels. Indeed, we have now reported a transition of a region of S6 to a *π* helix as a factor in opening of two bacterial channels, NavMs and NavAb. In addition, the opening of certain TRP channels appears to involve an α to *π* transition and a reorientation of a conserved Asn residue (31,69-72), raising the possibility that this is a conserved mechanism across evolution. Eukaryotic sodium channels also possess a conserved asparagine residue (N1211) in their S6 helices, and structures have revealed the presence of *π*-helices in certain states of some isoforms. Most human sodium channels feature a *π*-helix in domains I and II, while the other two S6 helices (domains III and IV) are α-helical (21,22,24-27). Recently, refined analysis of Nav1.7 densities showed a transient *π* helix in domain IV (33), indicating that this transition is indeed energetically possible. This conformational change has been linked to inactivation, posing provocative possibilities for future study of helical transitions in the gating cycle of both bacterial and eukaryotic channels. Another interesting consequence of *π*-helix formation in S6 was observed at the so-called fenestrations, which are thought to provide pathways for hydrophobic drugs such as lidocaine or benzocaine to access binding sites in Nav-channel pores (5). More precisely, hydrophobic drugs have been proposed to freely access the pore in closed states, but not in open ones (5). In NavAb we found that the fenestrations are closed in *π* models irrespective of the dimensions (constricted/expanded) of the pore, while they are open in α models. This finding is in agreement with a functional assignment of the eukaryotic channels NavPas and NALCN as pre-open (60,61,65), in which all S6 segments contain *π* helices and the fenestrations are obstructed. Moreover, recent structures of human Nav1.7 indicated that the presence of a *π* helix in subunit IV closed the fenestration at the interface with subunit I (33). Lidocaine binding to different functional states of NavAb revealed that the drug binds near the selectivity filter in resting and *π* models, in agreement with previous simulations in NavMs and NavPas, respectively (5, 66). Lidocaine binds nearer the intracellular gate in the expanded-α, presumed inactivated model. In the alternative, asymmetric inactivated model, lidocaine binds even further towards the intracellular space, forming contacts with V1213 in addition to T1206. Lidocaine interacts in all states with T1206, suggesting that this residue serves as an important contact for local anesthetics, in agreement with studies on NavAb (5) and NachBac (35) and on eukaryotic sodium channels (67, 68). In the eukaryotic sodium channel Nav1.2, lidocaine is known to bind to F1760 and Y1771, which align with T1206 and V1213 (67,68). This further highlights the importance of bacterial sodium channels as appropriate models to study the function and modulation of eukaryotic sodium channels.

## Acknowledgements

We acknowledge SciLifeLab and the Swedish Research Council to LD (VR 2018-04905) for funding. The MD simulations were performed on resources provided by the Swedish National Infrastructure for Computing (SNIC) at PDC Center for High Performance Computing (PDC-HPC) and the resources provided by Swiss national supercomputing center.

## Author contributions

K.C and L.D designed the research. K.C performed the simulations and analyzed the results. K.C, R.J.H and L.D interpreted the data and wrote the manuscript.

