## Supplementary Material for "An α-π transition in S6 shapes the conformational cycle of the bacterial sodium channel NavAb"

### Supplementary Figures

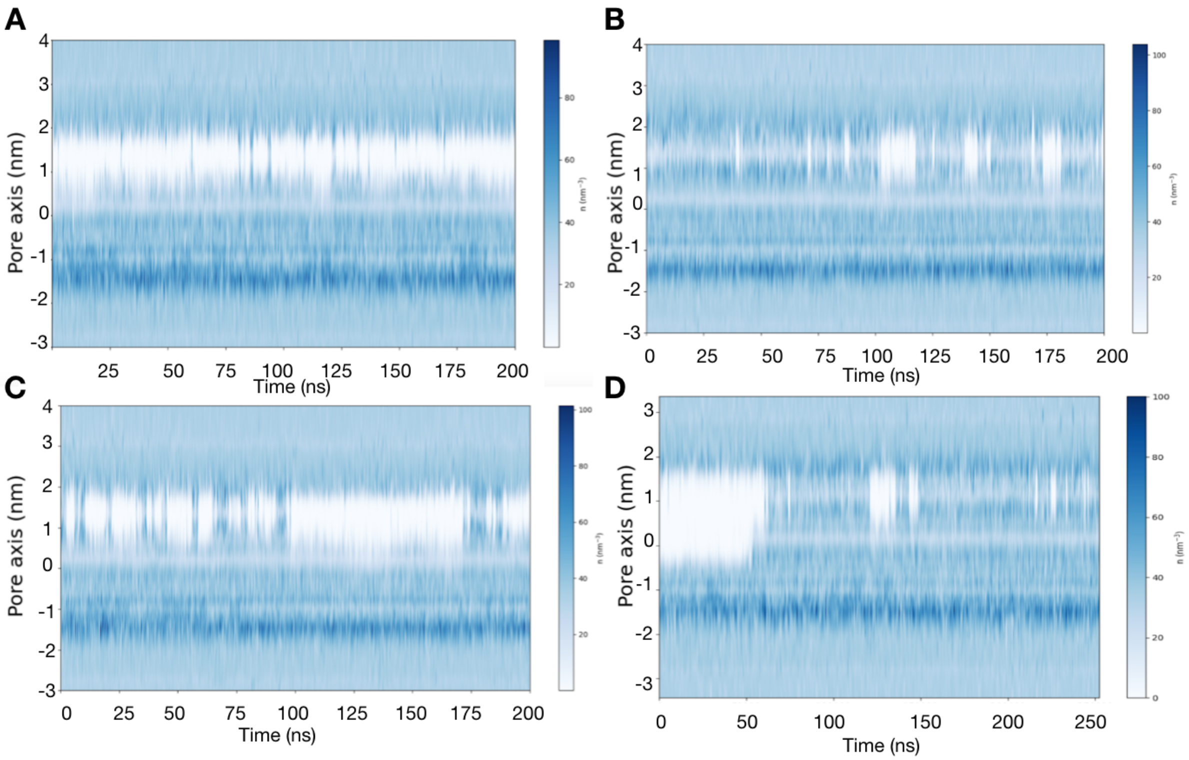


**Figure S1: A.** Water number density profile over time along the central pore axis of WT expanded-α model of NavAb at 0 mV. **B.** Water number density profile over time along the central pore axis at +750 mV. **C.** Water number density profile over time along the central pore axis at -750 mV. The activation gate is transiently hydrated **D.** Water number density profile over time along the central pore axis in the I1217C mutant at 0 mV.

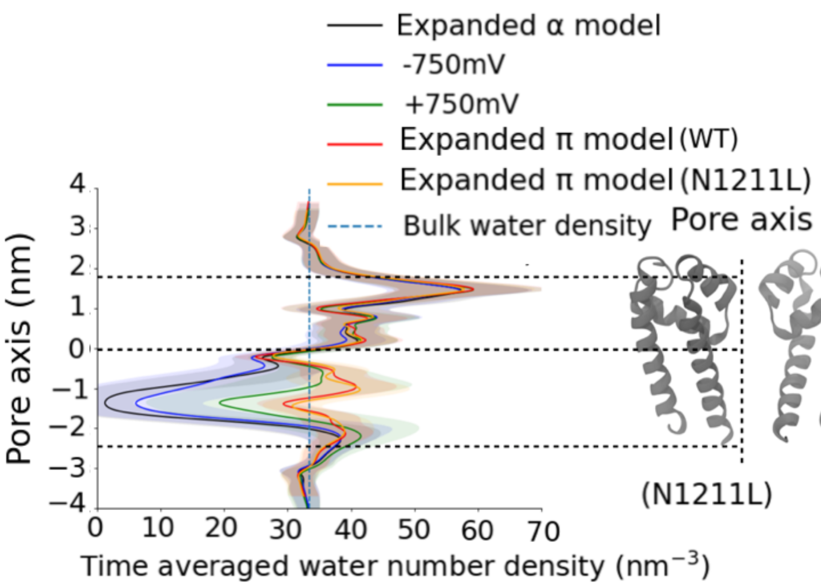


**Figure S2:** Time averaged water number density plot along the central pore axis for various NavAb models. Standard deviation shown in light shades of respective colors for different models.

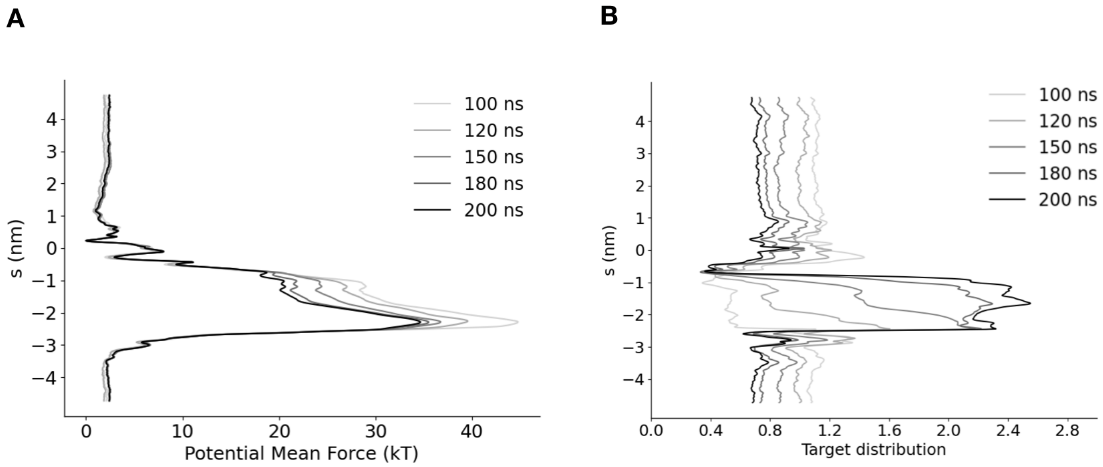


**Figure S3: A.** Convergence of free energy profile of sodium ion permeation in the NavAb expanded α model as determined using AWH. The free energy was calculated across 6 walkers sharing the bias. **B.** Target distribution at different times.

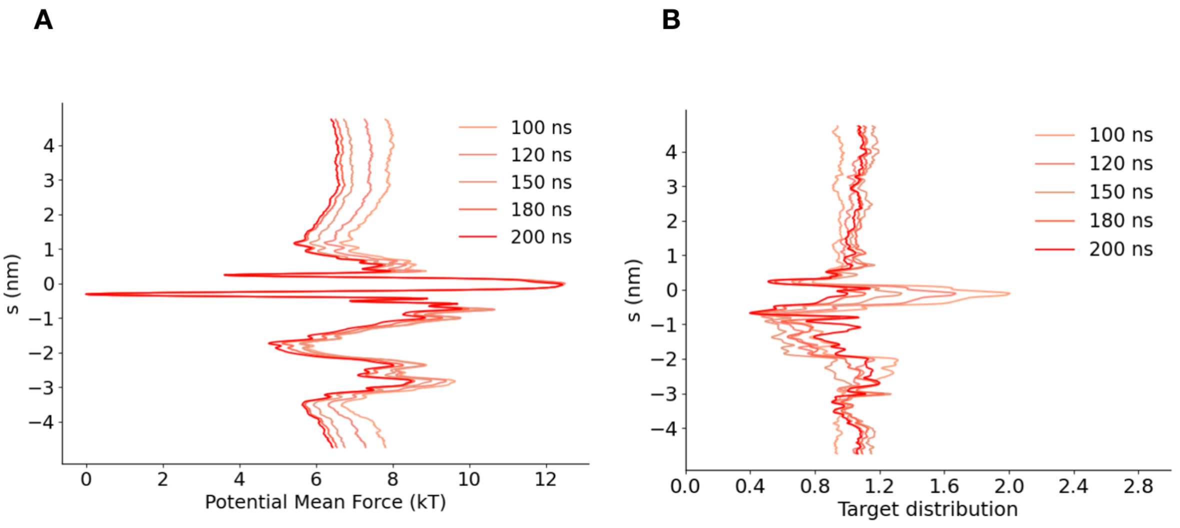


**Figure S4: A.** Convergence of free energy profile of sodium ion permeation in NavAb expanded $\pi$ model (WT) as determined using AWH. **B.** Target distribution at different times.

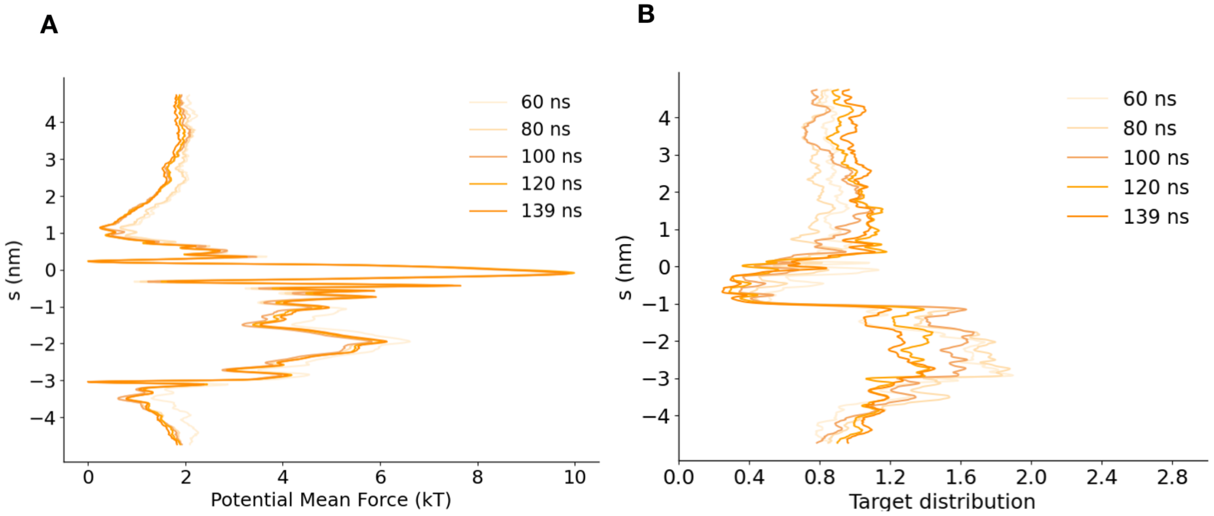


**Figure S5: A.** Convergence of free energy profile of sodium ion permeation in NavAb expanded $\pi$ model (N1211L) as determined using AWH. **B.** Target distribution at different times.

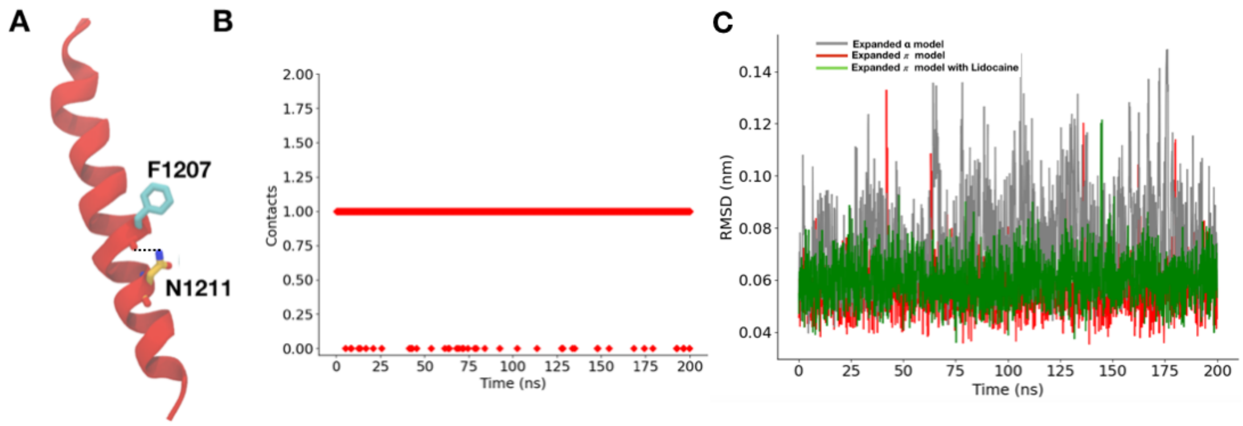


**Figure S6: A.** Backbone carbonyl of F1207 forming hydrogen bond with side-chain of N1211, hence stabilizing the $\pi$ helix. **B.** Number of hydrogen bonds between the F1207 backbone carbonyl and side-chain of N1211 in the $\pi$ helix model as a function of simulation time **C.** RMSD of N1211 in different models. Expanded α model - Gray, Expanded $\pi$ model - Red, Expanded $\pi$ model with Lidocaine - Green

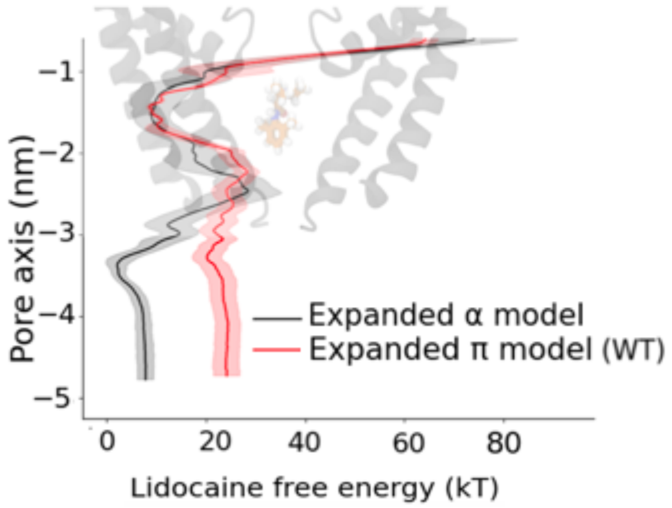


**Figure S7:** Lidocaine permeation free energy profiles as determined using AWH for the expanded α model (black) and expanded $\pi$ model WT (red) along the central pore axis of the lower half of the S6 helix. Cartoon representations in the background of D and E show the lower half of the S6 helix.

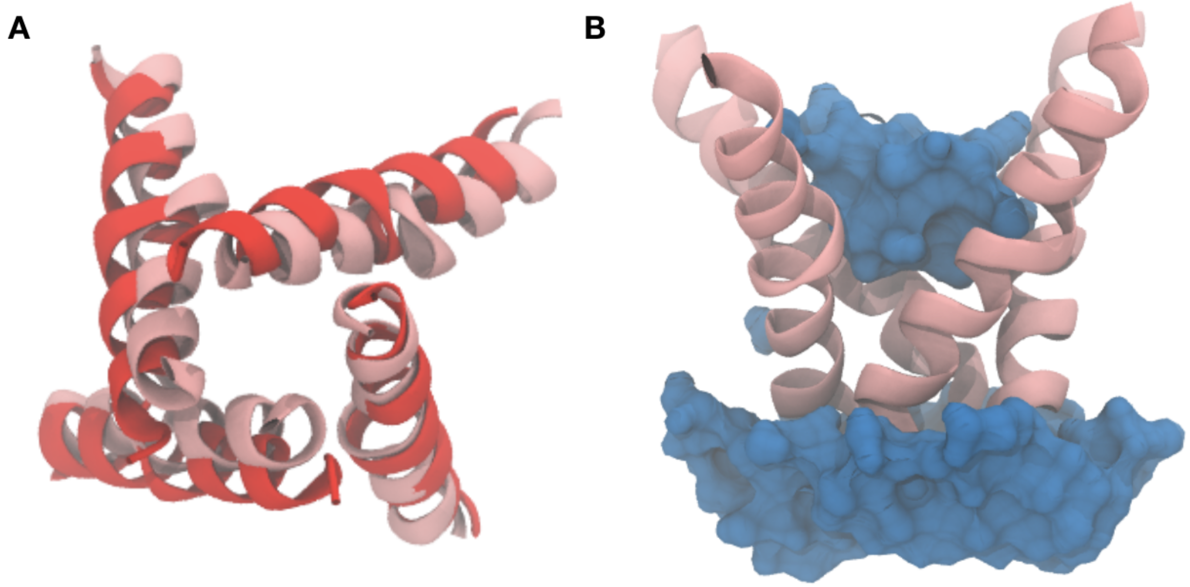


**Figure S8: A.** Structural alignment between the final model of the restrained simulations (red) and the unrestrained simulations (pink) shows that, in the absence of restraints, the pore collapses within 154 ns to a more contracted radius (pink), leading to dehydration (blue surface) around the constriction point (**B**).

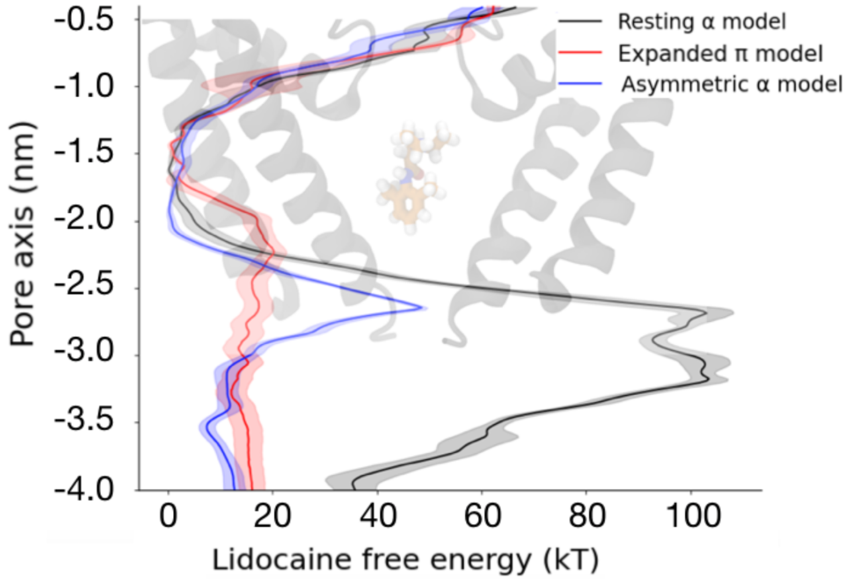


**Figure S9:** Lidocaine permeation free energy profiles as determined using AWH for the resting model (black), expanded $\pi$ model (red) and asymmetric α model (blue) along the central pore axis of the lower half of the S6 helix.

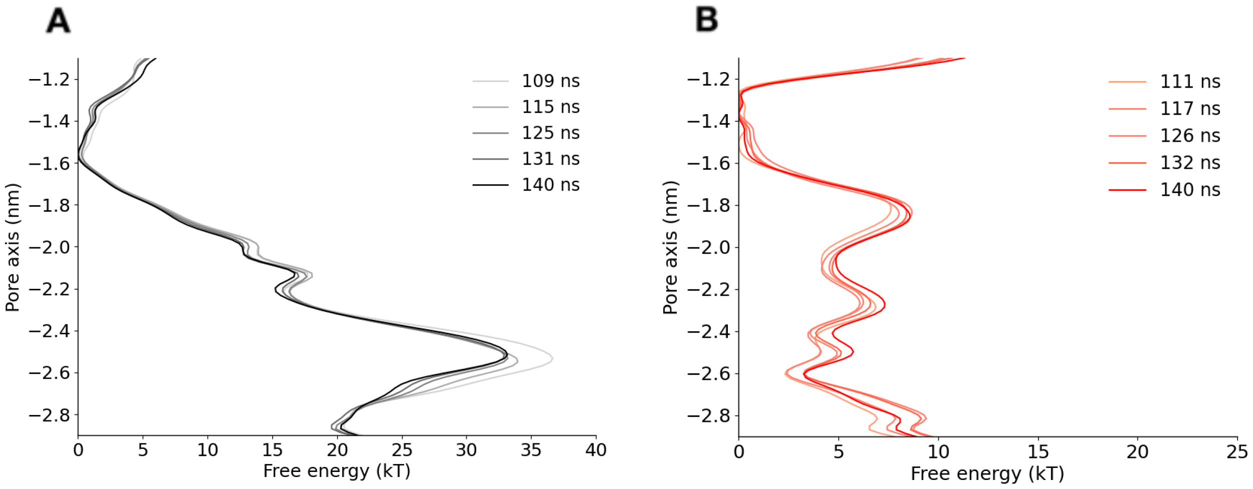


**Figure S10: A.** Convergence of free energy profile obtained using multiple walker well tempered metadynamics for lidocaine permeation in NavAb expanded α model. **B.** Convergence of free energy profile of Lidocaine permeation in NavAb expanded $\pi$ model.

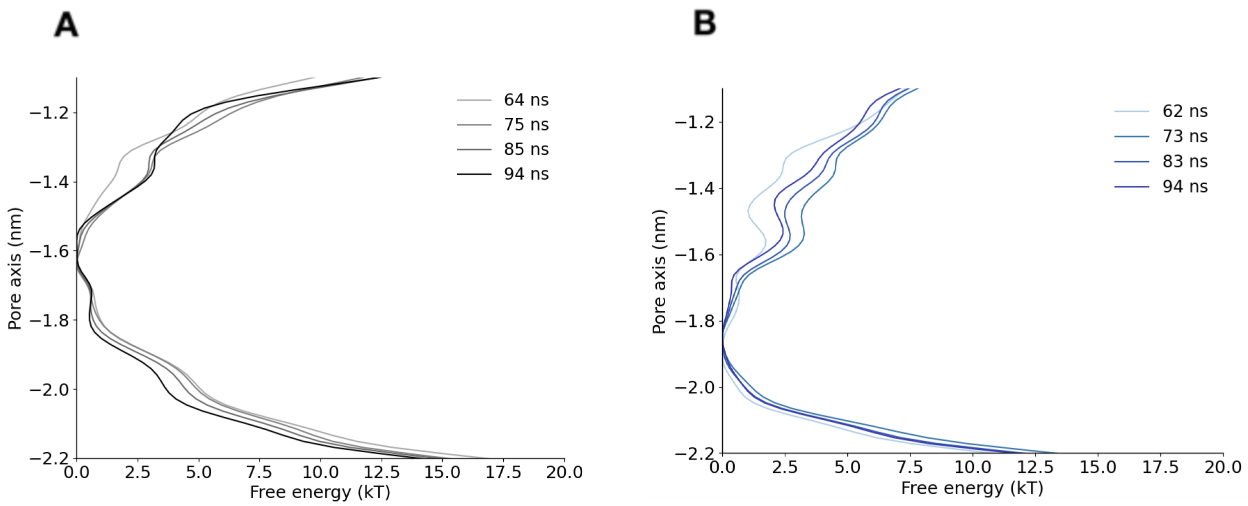


**Figure S11: A.** Convergence of free energy profile obtained using multiple walker well tempered metadynamics for Lidocaine permeation in NavAb resting model. **B.** Convergence of free energy profile of Lidocaine permeation in NavAb asymmetric α model.

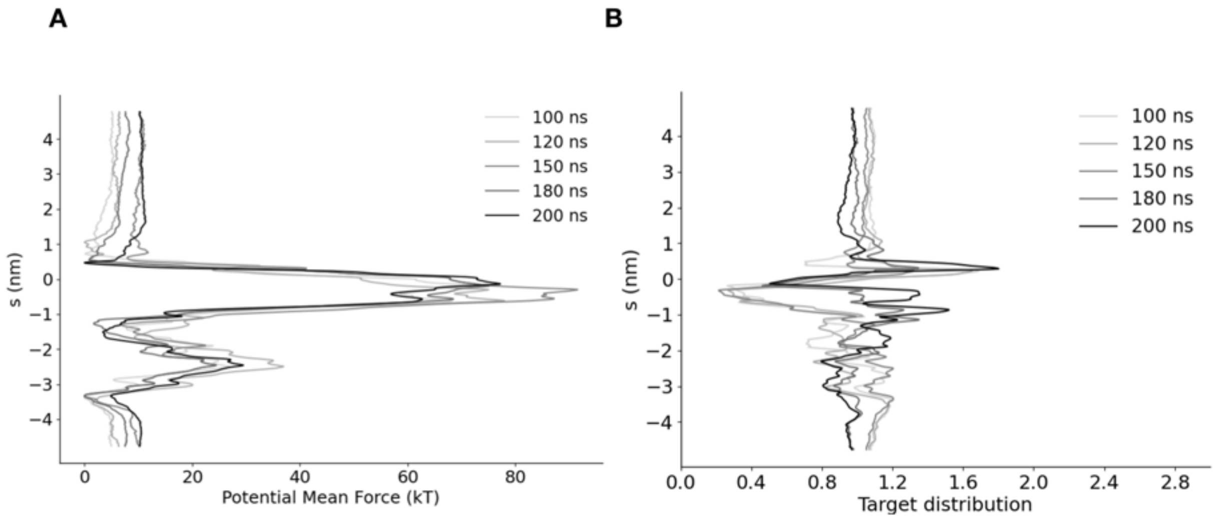


**Figure S12: A.** Convergence of free energy profile of lidocaine permeation in NavAb expanded α model as determined using AWH. **B.** Target distribution at different times.

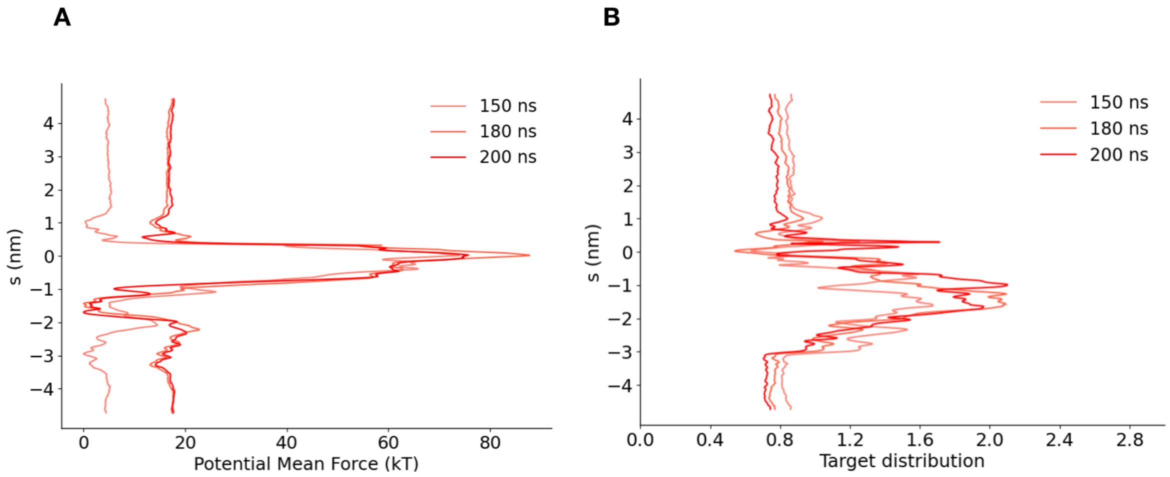


**Figure S13: A.** Convergence of free energy profile of Lidocaine permeation in NavAb expanded $\pi$ model as determined using AWH. **B.** Target distribution at different times.

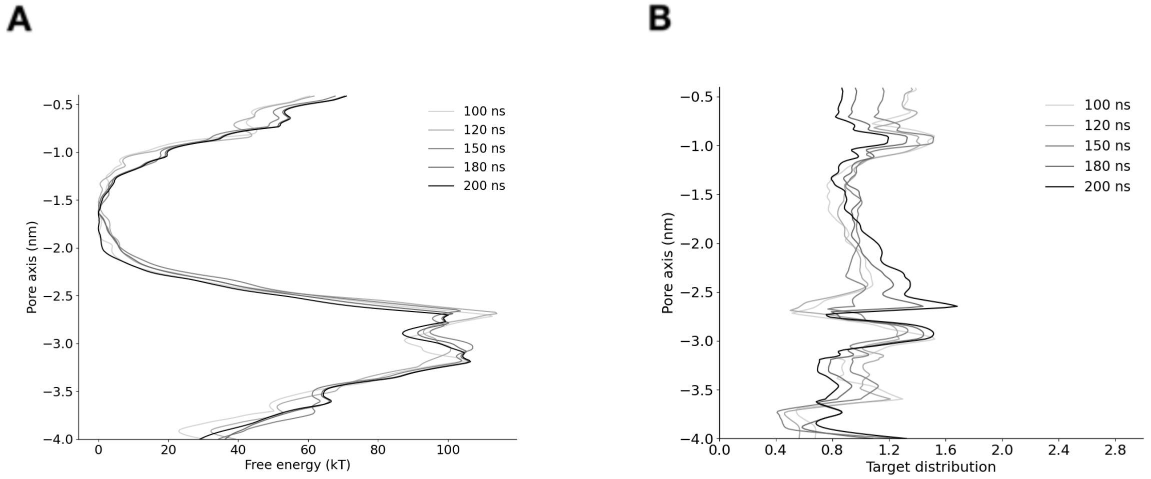


**Figure S14: A.** Convergence of free energy profile of Lidocaine permeation in NavAb resting model as determined using AWH. **B.** Target distribution at different times.

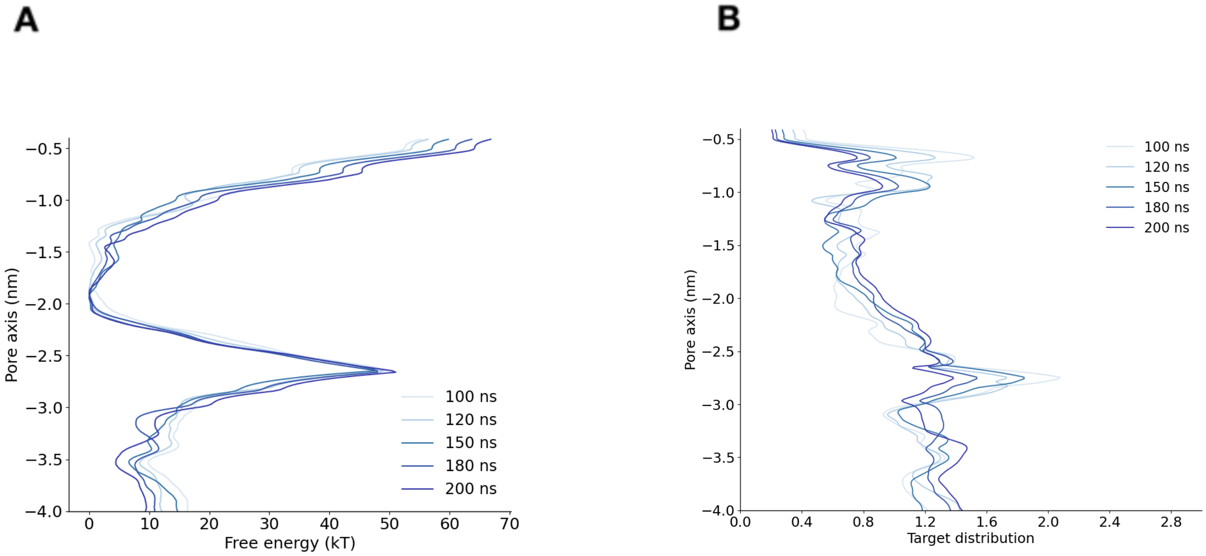


**Figure S15: A.** Convergence of free energy profile of Lidocaine permeation in NavAb asymmetric α model as determined using AWH. **B.** Target distribution at different times.

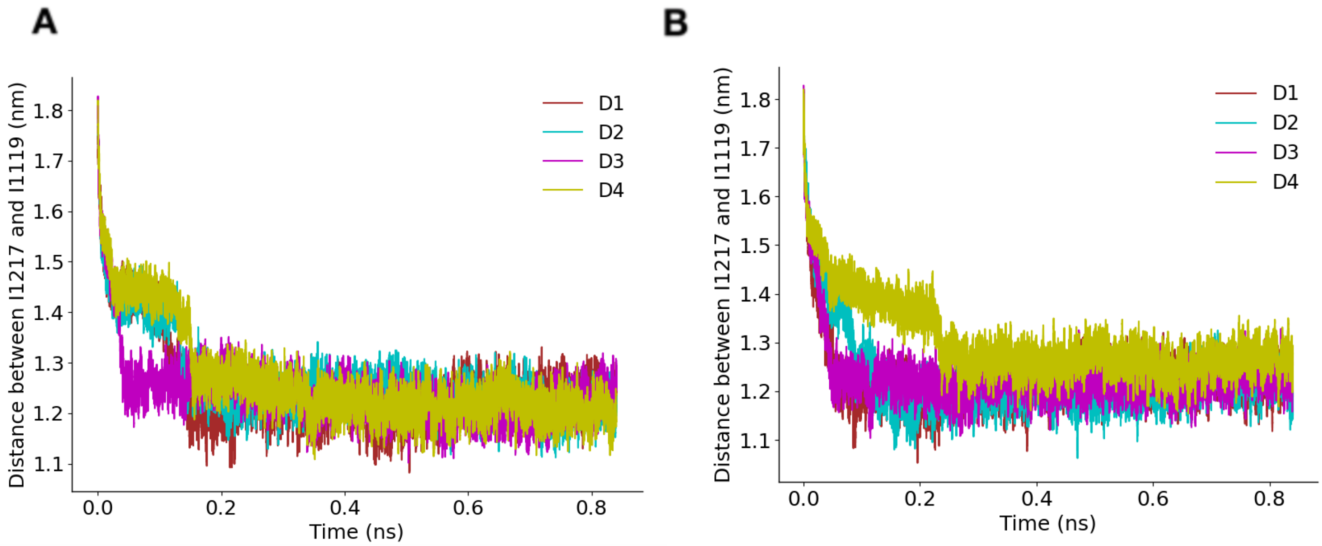


**Figure S16:** Time evolution of the pore opening ABMD simulation starting from the contracted $\pi$ model (WT) for two of the replicas (the other replica is found in Figure 5).

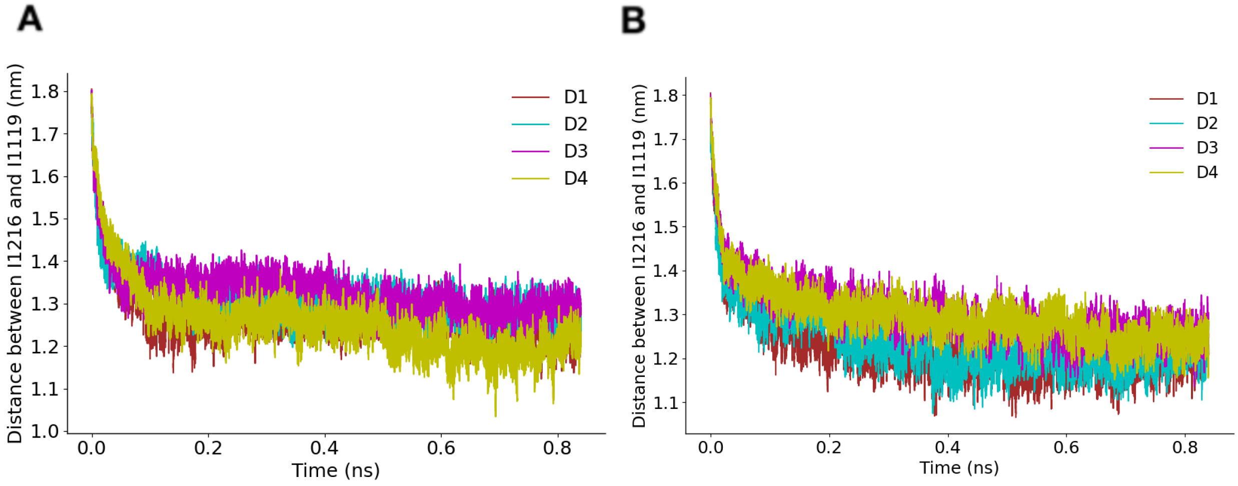


**Figure S17:** Time evolution of the pore opening ABMD simulation starting from the contracted α model for two of the replicas (the other replica is found in Figure 5).

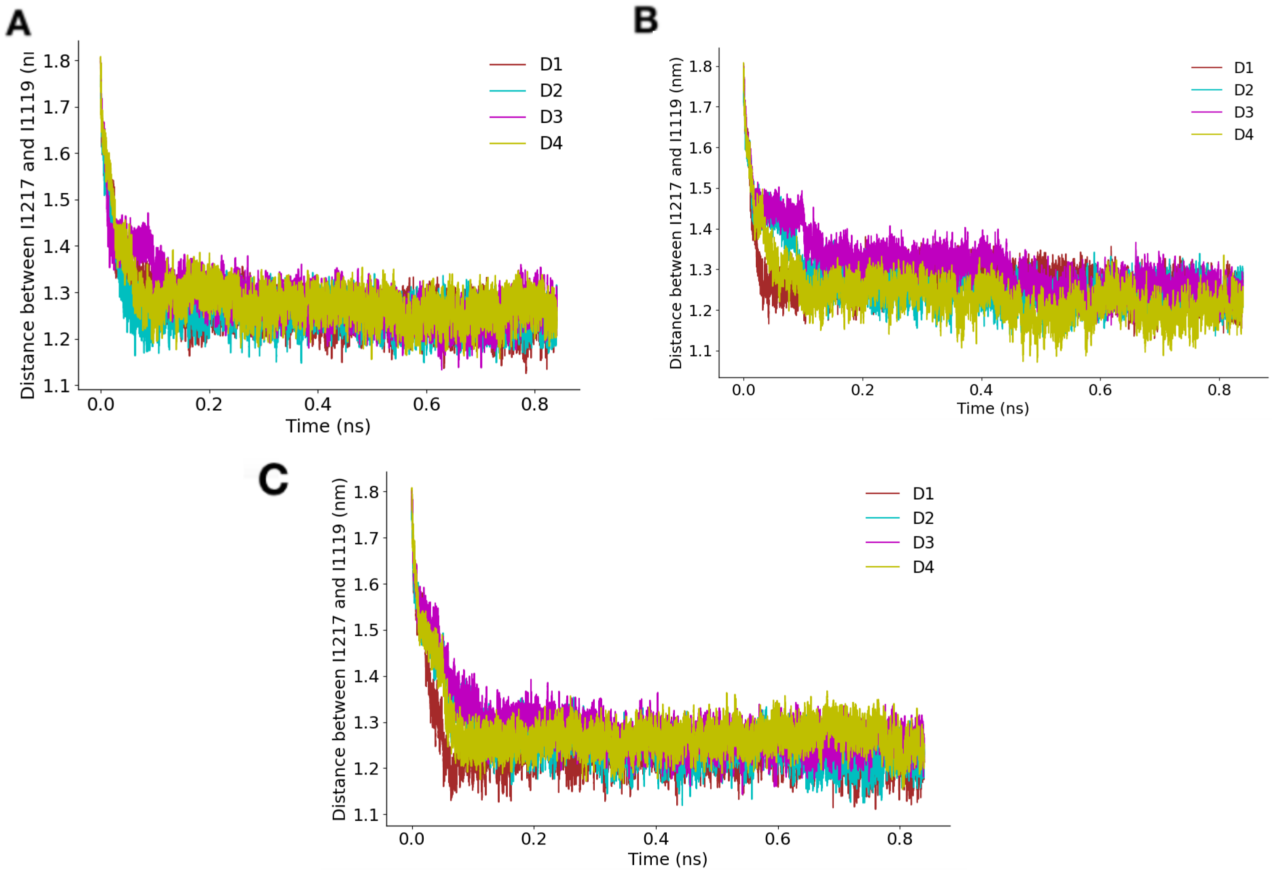


**Figure S18:** Time evolution of the pore opening ABMD simulation starting from the contracted $\pi$ model (N1211L) for three replicas.
